# Exploring the metabolic potential of *Aeromonas* to utilise the carbohydrate polymer chitin

**DOI:** 10.1101/2024.02.07.579344

**Authors:** Claudia G. Tugui, Dimitry Y. Sorokin, Wim Hijnen, Julia Wunderer, Kaatje Bout, Mark C.M. van Loosdrecht, Martin Pabst

**Affiliations:** Delft University of Technology, Department of Biotechnology, Delft, The Netherlands; Winogradsky Institute of Microbiology, Federal Research Centre of Biotechnology, RAS, Moscow, Russia; Evides Water Company, Rotterdam, The Netherlands

**Keywords:** *Aeromonas*, metabolism, secretomics, quantitative proteomics, chitin degradation

## Abstract

Members of the *Aeromonas* genus are commonly found in natural aquatic ecosystems. However, they are also frequently present in non-chlorinated drinking water distribution systems. High densities of these bacteria indicate favorable conditions for microbial regrowth, which is considered undesirable. Studies have indicated that the presence of *Aeromonas* is associated with loose deposits and the presence of invertebrates, specifically *Asellus aquaticus*. Therefore, a potential source of nutrients in these nutrient poor environments is chitin, the structural shell component in these invertebrates. In this study, we demonstrate the ability of two *Aeromonas* strains, commonly encountered in drinking water distribution systems, to effectively degrade and utilize chitin as a sole carbon and nitrogen source. We conducted a quantitative proteomics study on the cell biomass and secretome of both strains, revealing a dedicated and diverse spectrum of hydrolytic enzymes and pathways for the uptake and metabolism of chitin. Furthermore, when the primary nutrient source was switched from glucose to chitin, more than half of the *Aeromonas* proteome showed significant changes. Additionally, a genomic analysis of *Aeromonas* species found in drinking water distribution systems suggests a general potential ability of this genus to degrade and utilize a variety of carbohydrate biopolymers. This study indicates the relation between the utilization of chitin by *Aeromonas* and their association with invertebrates such as *A. aquaticus* in loose deposits in drinking water distribution systems. This knowledge provides the foundation for the development of more effective water sanitation strategies.

## Introduction

The genus *Aeromonas* comprises a group of Gammaproteobacteria that are widely distributed in aquatic environments. Some members of this genus have the potential to cause diseases in humans and other animals. *Aeromonas* are also commonly found in engineered ecosystems, such as drinking water distribution systems (DWDS). However, in these environments, the bacterium is generally considered non-pathogenic (1). Nevertheless, elevated levels of *Aeromonas* in non-chlorinated drinking water distribution systems (DWDS) are considered an indicator of favorable growth conditions for microbial growth, which may lead to compromised characteristics, including changes in taste, odour, colour, the presence of large invertebrates, as well as potential occurrence of opportunistic pathogens. Additionally, in the context of water resource management and global warming, maintaining the quality of drinking water is a persistent challenge. Therefore, studying the regrowth of microorganisms, such as *Aeromonas*, in DWDS is a crucial research objective.

The nutritional versatility of *Aeromonas* in the oligotrophic drinking water distribution systems has been investigated earlier, which showed a high affinity for amino acids, long-chain fatty acids (1, 2) and biopolymers such as starch and chitin (1, 3). However, experiments have shown that competitive planktonic growth in drinking water is not very likely (1). *Aeromonas* is commonly present in loose deposits and in bulk water of groundwater (4, 5) and surface water of drinking water distribution systems. This niche is also shared by invertebrates, including *A. aquaticus*. As a result, it has been speculated that *Aeromonas* may take advantage of the carbohydrate polymer chitin, which is a key component of the exoskeletons of these invertebrates.

Chitin is generally considered the second most abundant biopolymer in nature and is the most prevalent in aquatic environments such as the oceans. As a result, this polymer plays a crucial role in the global carbon and nitrogen cycles (6). The ability to utilize this carbohydrate polymer as nutrient source is therefore widely found among microorganisms (7-9). However, its role in environments such as the drinking water distribution systems is only poorly understood to date.

The utilization of this carbohydrate polymer is known to require a cascade of enzymatic reactions to make it accessible for uptake and growth. The chitin backbone is made of β-(1-4)-linked N-acetylglucosamine (GlcNAc) units. The hydrolytic degradation of chitin usually starts outside the cell via different (endo- and exo-) chitinases and associated hydrolytic enzymes. Additional lytic polysaccharide monooxygenases (LPMOs) can cleave crystalline chitin via an oxidation reaction. This generates regions with decreased crystallinity which subsequently become more accessible for other chitinases (10). Endochitinases cleave the chitin chain at random points to produce oligomers such as chitotetriose, chitotriose and chitobiose. Exochitinases and N-acetylhexoseamindases produce the smaller chitobiose and GlcNAc monomers (6). Chitin oligomers may also be deacetylated, which can be converted by chitosanases and hexosaminidases into glucosamine forms, and which may be even further cleaved by (non-specific) cellulases (6, 11, 12). The uptake of the GlcNAc oligomers and monomers can be facilitated through dedicated porins, ABC and PTS transporters (13). Further catabolism of the GlcNAc in the cytoplasm commonly starts with phosphorylation by a GlcNAc kinase (6). Alternatively, PTS sugar transporters perform transport and phosphorylation simultaneously (6, 14). The resulting GlcNAc-P can then be deacetylated and deaminated to produce fructose-6P, a metabolite which can directly enter glycolysis (6). Additionally, the conversion of GlcNAc into GlcNH2 releases acetate, which further results in the production of acetyl-CoA, and the deamination releases ammonia, which can be incorporated into the proteinogenic amino acid glutamine (6). Glutamine furthermore plays an important role as a nitrogen carrier and storage within bacterial cells (15). Finally, the oxidative degradation of chitin by LPMOs produces oxidized GlcNAc mono- and oligomers. These oxidation products, have been reported to be directly deacetylated and deaminated to produce 2-keto-3-deoxygluconate, acetate and ammonia without prior phosphorylation (16). This is supposed to be a common utilization route for crystalline chitin in marine Gammaproteobacteria (16). Several bacteria, including some *Aeromonas* strains, have already been investigated for the expression and activity of a range of chitinases, such as *A. hydrophila* (14), *Aeromonas sp*. (17) and *Aeromonas caviae* (18). Furthermore, the suspended growth of some *Aeromonas* strains on low concentrations of chitin and nitrate was demonstrated recently (1).

However, the underlaying metabolic routes and the cellular and secretome response to chitin as sole carbon and nitrogen source, has not been investigated to date. Therefore, we demonstrate the efficient growth on chitin as sole carbon and nitrogen source for two selected *Aeromonas* species, *Aeromonas bestiarum* and *Aeromonas rivuli*, which are frequently found in non-chlorinated drinking water distribution systems (1). We performed an extensive quantitative proteomics study on their chitin degradation, uptake and catabolic network. Remarkably, the switch from glucose to chitin induced changes in 65% of the quantified proteins in both strains. The identified upregulated secreted enzymes, porins and transporters provide strong evidence for a dedicated chitin utilization network in *Aeromonas* members. Finally, a broader study on biopolymer degradation routes suggests that the genus *Aeromonas* possesses the ability to break down a range of different biopolymers. In summary, the study brings fundamental insights into the metabolic ability of *Aeromonas* in degrading and utilizing the carbohydrate polymer chitin as sole carbon and nitrogen source. This provides a better understanding of how these microbes can survive in nutrient-poor environments such as drinking water distribution systems.

## Materials and methods

### Growth of *Aeromonas* on glucose and chitin

Two *Aeromonas* strains (*A. bestiarum* DSM 13956 and *A. rivuli* DSM 22539) were purchased from DSMZ (Leibniz, Germany, https://www.dsmz.de/dsmz). The cells were reactivated, and grown on glucose and peptone rich TSB medium, as recommended by DSMZ. From these cultures, glycerol (30%) stocks were obtained which were stored at -80°C until further use. The two *Aeromonas* strains, *A. rivuli* and *A. bestiarum* were inoculated on M9 minimum salt medium (Na2HPO4 6.78 g/L, KH2PO4 3 g/L, NaCl 0.5 g/L, NH4Cl 0.21 g/L) and acidic trace element solution (19) combined with 20% MgSO4 1:1 to (2 mL to 1 L of medium), and 10% CaCl2 (1 mL/L of medium), pH=6.7. Glucose was added to a concentration of 13.6 mM, and amorphous chitin, prepared through ‘HCl decrystallization’ from powdered defatted/deproteinated crab shells (Sigma Aldrich/Merck, US)(20), was added to a final concentration of 0.5 g/L. The cultures were grown aerobically at 33°C on a rotary shaker (Edmund Bühler GmbH, Germany) at 100 rpm. The growth was monitored using an Ultrospec 10 Cell Density Meter (Biochrom Ltd, UK). For both strains we conducted biological triplicates of both growth conditions (glucose and chitin), resulting in a total of 12 shake-flask experiments. **Microscopy**. Light microscopy was performed using a Zeiss Axio Imager M2 microscope equipped with an Axiocam 305 color camera (Carl Zeiss, Germany). The microscope setting possesses a 63x and 100x oil immersion objective lens and phase contrast capabilities. The proprietary Zeiss software for image capture and analysis was Zen 3.3. **Cell harvesting and supernatant concentration**. From every culture, every day 1 mL of cell broth was harvested and centrifuged (14K rpm, 10 minutes) to separate the cell pellet from the supernatant. The resulting supernatant (approx. 1 mL) and cell pellet was stored separately at -20°C, until further processed. For the analysis of chitin hydrolysis products, 50 mL of the supernatant was further filtered through sterile syringe filters (0.2 μm Sartorius) and then concentrated using a speedvac concentrator to a final volume of 500 μL. The time point with the highest OD was selected for proteome analysis. **Cell lysis, protein extraction, and proteolytic digestion**. *Analysis of the cell biomass proteome*. The cell pellets from the biological triplicates of each strain (*A. bestiarum* and *A. rivuli*) and contrasted growth conditions (glucose and chitin) were resuspended in 175 μL 50 mM TEAB buffer (with 1% NaDOC) and 175 μL B-PER buffer (Thermo Scientific, Germany) by vortexing. Then acid washed glass beads (105– 212μm, Sigma Aldrich), were added and the mixtures were vortexed thoroughly, sonicated for 15 minutes and then frozen at -80°C for 15 minutes. Thereafter, the samples were thawed in a Thermocycler at 40°C and under shaking at 1000 rpm for 15 minutes. Afterwards, the samples were spun down at 14000 rpm. The supernatant was collected, and TCA was added (1 volume TCA to 4 volumes supernatant). The mixture was vortexed and incubated at 4°C for 20 minutes, then spun down at 14000 rpm for 15 minutes at 4°C. The obtained protein pellets were once washed with ice cold acetone and then dissolved in 6 M urea (in 100 mM ammonium bicarbonate, ABC). Further, the disulfide bridges were reduced by the addition of 10 mM DTT and incubation for one hour at 37°C under shaking at 300 rpm. Thereafter, 20 mM IAA was added. The mixture was kept in the dark for 30 minutes. 200 mM ABC buffer was then added to the samples to obtain a solution with <1 M Urea. Finally, proteolytic digestion was performed by adding Trypsin (0.1 μg/μL in 1 mM HCl, Sequencing Grade Modified Trypsin, Promega) at a ratio of 50:1 (w:w, Protein:Trypsin) to the sample. The proteolytic digestion was performed overnight at 37°C, under gentle shaking at 300 rpm. Peptides were desalted using an OASIS HLB solid phase extraction well plate (Waters, UK) according to the instructions of the manufacturer, speed vac dried and stored at -20°C until further processed. *Analysis of the secreted proteome*. 600 μL of the collected supernatants from the biological triplicates of each strain (*A. bestiarum* and *A. rivuli*) and both growth conditions (glucose and chitin) were processed with the same protocol as described for the cell pellets (see above) albeit starting directly with the TCA protein precipitation step. **Quantitative shotgun proteomics and analysis of the secreted proteome**. Approx. 500 ng of proteolytic digest were analyzed in duplicate injections using an EASY nano-LC 1200, equipped with an Acclaim PepMap RSLC RP C18 separation column (50 μm x 150 mm, 2 μm), and a QE plus Orbitrap mass spectrometer (Thermo Fisher Scientific, Germany). The flow rate was maintained at 350 nL/min over a linear gradient from 5% to 25% solvent B over 180 min, then from 25% to 55% B over 60 min, followed by back equilibration to starting conditions. Data were acquired from 5 to 240 minutes. Solvent A was H2O containing 0.1% formic acid, and solvent B consisted of 80% ACN in H2O and 0.1% formic acid. The mass spectrometer was operated in data-dependent acquisition mode. Full MS scans were acquired from m/z 380–1250 at a mass resolution of 70 K with a maximum injection time (IT) of 75 ms and an automatic gain control (AGC) target of 3E6. The top 10 most intense precursor ions were selected for fragmentation using higher-energy collisional dissociation (HCD). MS/MS scans were acquired at a resolution of 17.5 K with an AGC target of 2E5 and IT of 75 ms, 2.0 m/z isolation width and normalized collision energy (NCE) of 28.

### Processing of mass spectrometric raw data

Database searching of the shotgun proteomics raw data was performed using proteome reference databases from *A. bestiarum* and *A. rivuli*, obtained from UniprotKB (UP000224937) and NCBI (NCBI taxonomy ID: 648794), using PEAKS Studio X (Bioinformatics Solutions Inc., Canada). The database searching allowed 20 ppm parent ion and 0.02 m/z fragment ion mass error, 3 missed cleavages, carbamidomethylation as fixed and methionine oxidation and N/Q deamidation as variable modifications. Peptide spectrum matches were filtered for 1% false discovery rates (FDR) and identifications with ≥2 unique peptides were considered as significant. Quantitative analysis of the changes between chitin and glucose-grown conditions, and the cell pellet and secreted proteome abundances was performed using the PEAKSQ module (Bioinformatics Solutions Inc., Canada). Normalization was based on the total ion current (TIC), and only proteins with at least 2 unique peptides and identified in at least 2 out of 3 biological replicates were considered. Peptide spectrum matches were filtered with a 1% false discovery rate (FDR). ANOVA was used to determine the statistical significance of the changes between the conditions, expressed as -10xlog10(p), where p corresponds to the significance testing p-value. **Annotation of structural components, functions and estimation of protein abundance**. Results from PEAKSQ were further used for the analysis of enriched functions and metabolic pathways. Data processing and visualization was performed using Python 3.11.3. Furthermore, SignalP 6.0 (https://services.healthtech.dtu.dk/services/SignalP-6.0/) (21) was used for the prediction of signal peptides in order to confirm secreted proteins (prediction > 0.9 was required to accept Sec or TAT annotation). Proteomes for both *Aeromonas* species were uploaded to STRING (version 12.0, https://string-db.org/) to generate interaction networks and to annotate with GO terms (*A. bestiarum* string taxid: STRG0A90GTX, *A. rivuli* string taxid: STRG0A19KWE). The STRING tool was furthermore used to identify the enriched GO terms between conditions using proteins with >1.5-fold or <0.66-fold change. The HMM 3.3.2 tool (http://hmmer.org/) was used with the Pfam (http://pfam-legacy.xfam.org) (22) and the Carbohydrate Active Enzymes (CAZy) (http://www.cazy.org/) (23) databases to identify the glycolytic enzymes and their associated modules. For this, default parameters were used to perform an HMM scan. The results were filtered for e-values <10E-5 and for independent e-values of <10E-5. The transporters were identified by using a regex search using the terms: “PTS”, “porin”, “permease”, “transporter”, “transport”. Different ABC transporters and their subunits were grouped into general “ABC transporter” categories in the figures. The reference proteomes were furthermore annotated with KO numbers using BlastKoala (https://www.kegg.jp/blastkoala/), and the KEGG pathway map was obtained from Kegg-mapper (https://www.kegg.jp/kegg/mapper/) using the annotated proteins. emPAI indices were calculated for the central carbon metabolism enzymes according to the formula: emPAI=10^(#observed/#observable)-1, where *Nobserved* is the number of peptides measured in the experiment and *Nobservable* is the number of theoretical peptides that a protein can produce (24). The considered mass range for theoretical peptides was 800–3000 Da. For the multisubunit enzymes, the ratio glucose/chitin was determined by first averaging the area of the subunits and then the ratio was determined by diving the area of the respective enzymes. For homologue enzymes, only the variant with the highest sequence coverage is shown in the figures. The complete list of identified enzymes for each metabolic pathway is provided in the SI-doc (SI Table 1). The analysis for potential other carbohydrate polymer degrading genes of different *Aeromonas* strains was performed using the HMM 3.3.2 tool (http://hmmer.org/) and the Carbohydrate Active Enzymes (CAZy) database (http://www.cazy.org/) as described above. For this, the following proteomes were obtained from UniProt: *Aeromonas hydrophyla* (TaxID 644, UP000000756), Aeromonas *media* (TaxID 651, UP000502657), *Aeromonas caviae* (TaxID 648, UP000280168), *Aeromonas encheleia* (TaxID 73010, UP000275277), *Aeromonas molluscorum* (TaxID 271417, UP000013526), *Aeromonas salmonicida* (TaxID 645, UP000077360), *Aeromonas schubertii* (TaxID 652, UP000058114), *Aeromonas taiwanensis* (TaxID 633417, UP000297311), *Aeromonas veronii* (TaxID 654, UP000237142). The relevant CAZy database families for the different carbohydrate polymers are shown in the Supplementary Information Table 1.

### Chitin degradation assay

30 μL unfiltered supernatant of *A. bestiarum* and *A. rivuli* cultures were incubated with 1 mg of chitin suspended in 220 μL 1% PBS (phosphate buffered saline, Sigma Aldrich/Merck, US) in LC-MS grade water. Additional control samples were prepared containing only supernatant in 1% PBS, or chitin in 1% PBS. The samples were incubated at 33°C, 300 rpm for 18.5 hours. The samples were further spun down at 10K rpm using a bench top centrifuge for 2.5 minutes. GlcNAc oligomer standards were prepared from chitin following hydrolysis using HCl. For this, 250 μL of 5 M HCl was mixed with 1 mg of chitin. The mixture was incubated at 37°C, under shaking at 300 rpm for 18 hours. The sample was centrifuged at 14K rpm for 10 minutes. The supernatant was then purified before MS analysis using 25 mg HyperSep™ Hypercarb™ solid phase extraction cartridges (Thermo Fisher Scientific, Germany). The PGC material was washed with 500 μL 50% acetonitrile (0.1% formic acid), and then equilibrated with 2 x 500 μL H2O. samples were then loaded on the PGC cartridge and washed with 1 x 500 μL H2O. Sugars were eluted with 1 x 300 μL 50% acetonitrile (0.1% formic acid), collected in an Eppendorf tube and speed-vac dried. Samples were stored at -20°C until further analyzed. **MS analysis of chitin hydrolysis products**. SPE purified samples from the release assays were analyzed using an ACQUITY UPLC system (Waters, UK) equipped with a Hypercarb™ separation column (100 x 1 mm, Thermo Scientific, Germany) which was connected to a QE Focus™ hybrid quadrupole-Orbitrap mass spectrometer (Thermo Fisher Scientific, Germany). Solvent A was 100% water (0.1% formic acid) and solvent B was 100% acetonitrile (0.1% formic acid). A gradient was maintained at 100 μL/min flow rate from 2.5% B to 40% B over 8 minutes, and constant 40% B over 5 minutes, before equilibrating back to 2.5% B. The mass spectrometer was operated in ES+ (3.25 kV), where full MS scans were acquired from 250–1500 m/z at 70K resolution and an AGC target of 1e6. The m/z values for the GlcNAc mono and oligomers are GlcNAc= C8H16NO6+, 222.09721; GlcNAc-GlcNAc= C16H29N2O11+, 425.17659; GlcNAc-GlcNAc-GlcNAc= C24H43N3O16+, 629.26378; and for the oxidized forms GlcNAc1A= C8H16NO7+, 238.09213; GlcNAc-GlcNAc1A= C16H29N2O12+, 441.1715, and for the native deacetylated forms GlcNAc-GlcNH2 = C14H27N2O10+, 383.16602. Additional parallel reaction monitoring (PRM) was performed for the native GlcNAc mono and oligomers: m/z 222 ([M+H]+, HexNAc), m/z 425 ([M+H]+, (HexNAc)2), m/z 629 ([M+H]+, (HexNAc)3), m/z 833 ([M+H]+, (HexNAc)4), and m/z 1033 ([M+H]+, (HexNAc)5) using an isolation window of 2 m/z, an AGC target of 1e5, 100 ms max IT, 2 micro scans and 35K resolution. MS raw data were analyzed using XCalibur 4.1, where the area for the MS1 precursor ion or the most abundant fragment ion for each compound was integrated. **Data availability**. The shotgun proteomics raw data, reference sequence databases and database searching files have been deposited in the ProteomeXchange consortium database with the dataset identifier PXD047459.

## Results

### A quantitative proteomics study on *Aeromonas* grown on glucose and chitin

Two *Aeromonas* strains, *A. bestiarum* and *A. rivuli* were cultured in the presence of either glucose or amorphous chitin and subjected to a quantitative proteomics study. The cell culture supernatants were furthermore subjected to a chitin degradation study in order to identify the size distribution of the chitin degradation products (Figure 1). Culturing experiments were performed in biological triplicates for both strains, resulting in a total of 12 shake flask experiments. Both *Aeromonas* strains showed immediate growth on chitin (SI Figure 1) Nevertheless, microscopy images showed for both strains homogeneous cultures with rod-shaped cells approximately 1–2 μm in length (SI Figure 2).

**Figure 1:**
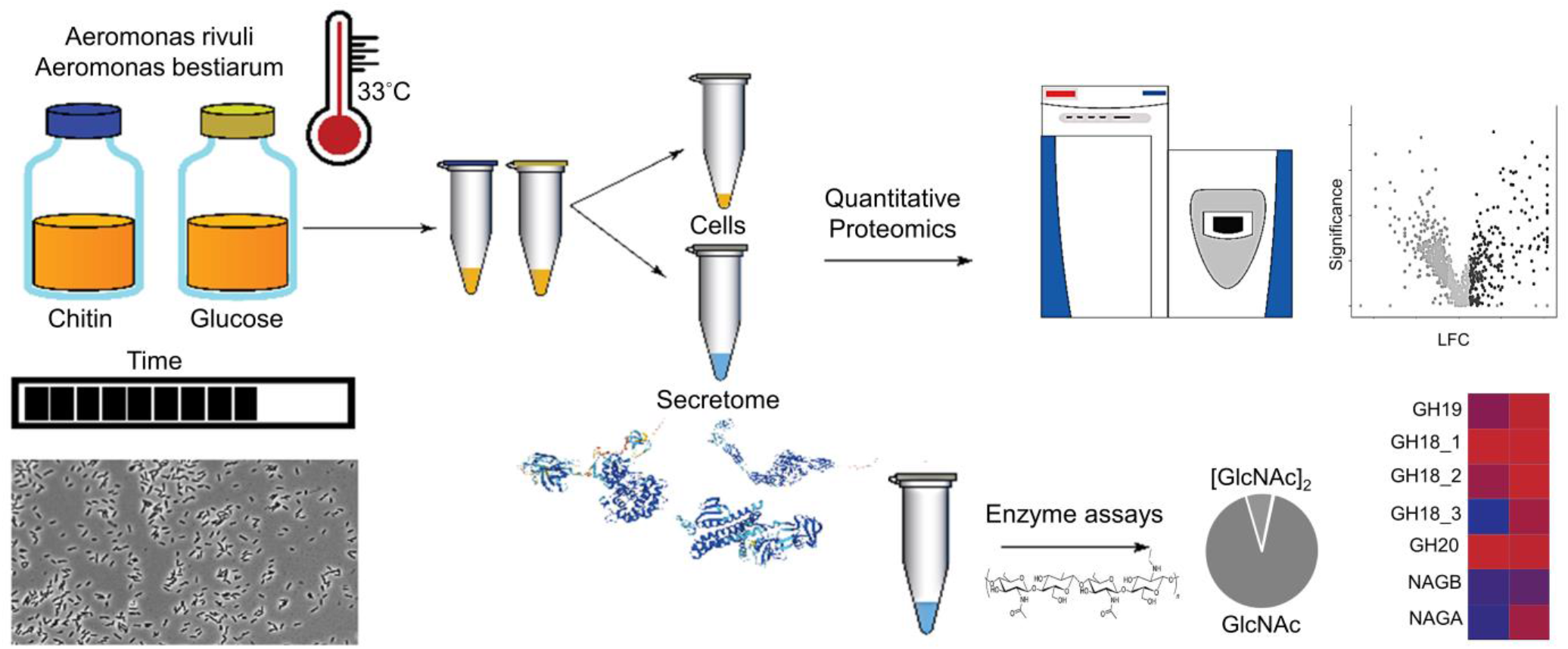
The graph outlines the employed workflow used to study the chitin degradation and utilization routes in *A. bestiarum* and *A. rivuli*. Both *Aeromonas* strains were cultured at 33°C with either glucose or chitin in biological triplicates. The growth of the bacteria was followed using OD660 and light microscopy over time. Quantitative proteomics was then employed to identify the enzymes secreted by the bacteria and to reveal their uptake and metabolic routes when switching the growth substrate from glucose to chitin. Annotation of hydrolytic enzymes was furthermore performed using the CAZy reference database, and enriched and depleted functions and pathways were identified using the STRING tool. Finally, chitin degradation assays were performed using the supernatants of both strains to determine the size distribution of the chitin hydrolysis products.

**Figure 2:**
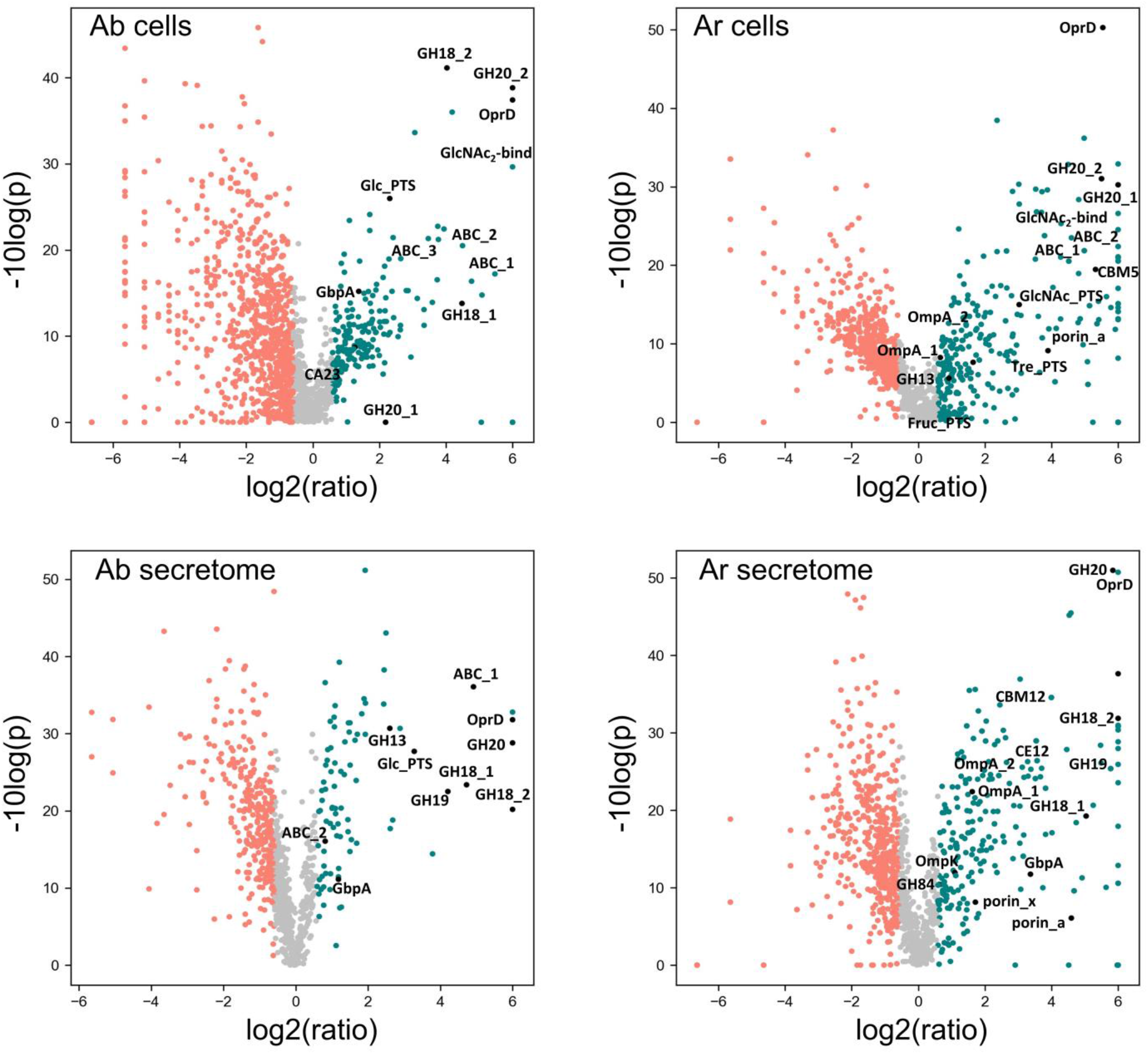
The volcano plots show the downregulated (red) and overexpressed (green) proteins in *A. bestiarum* (‘Ab’ left plot) and *A. rivuli* (‘Ar’ right plot) when comparing growth on glucose with cells grown on chitin. The x-axis shows the log2(fold-change) of the ratio chitin/glucose, and the y-axis shows the -10log(p) value of each fold-change, where p represents the statistical significance of the fold change. The upper plots show the cell biomass (‘cells’) proteome changes, where the lower plots show the secretome proteome changes. *A. rivuli* (Ar) cells: GH20_1 WP_224432421, GH20_2 WP_042039723, GH13 WP_224431530, CBM5 WP_232301920, OprD WP_042041362, OmpA_1 WP_224430968, porin_a WP_042043439, OmpA_2 WP_042039866, GlcNAc_PTS WP_042039724, Tre_PTS WP_042041746, Fruc_PTS WP_042039507, GlcNAc2-bind WP_042041499, ABC_1 WP_042041497, ABC_2 WP_224432441. *A. rivuli* (Ar) secretome: GH18_1 WP_042040344, GH18_2 WP_224432263, GH19 WP_084218236, CBM12 WP_232301920, GH20 WP_224432421, GH84 WP_224432541, GbpA WP_232302089, CE12 WP_042041941, porin_x WP_042044143, OmpA_1 WP_042040660, OmpK WP_224431060, OprD WP_042041362, OmpA_2 WP_042040662, porin_a WP_042043439. *A. bestiarum* (Ab) cells: GH18_1 A291U6Z3, GH18_2 A0A291TVA9, GH20_1 A0A291U0V8, GH20_2 A0A291U507, OprD A0A291U719, GbpA A0A291TWD8, CA23 A0A291U526, Glc_PTS A0A291U5N4, GlcNAc2-bind A0A291U0Y6, ABC_1 A0A291TZJ5, ABC_2 A0A291TWX6, ABC_3 A0A291U6T2. *A. bestiarum* (Ab) cells: GH18_1 A0A291TVA9, GH18_2 A0A291U6Z3, GH19 A0A291U6T4, GH20 A0A291U507, GH13 A0A291U830, GbpA A0A291TWD8, OprD A0A291U719, ABC_1 A0A291U0Y6, ABC_2 A0A291U6T2, Glc_PTS A0A291U5N4.

Samples were selected from the plateau of the growth curves (between days 2–3) for subsequent proteomics and chitin degradation experiments. After centrifugation of the cell suspension, the cell pellet was separated from the supernatant and processed separately. All 24 samples (2 strains, 3 biological replicates for growth on glucose and chitin, and for each condition biomass and supernatant samples) were subjected to shotgun proteomic experiments. The reproducibility of the biological experiments was further confirmed by principal component analysis and hierarchical clustering of the obtained proteomics profiles (SI Figures 3 and 4). Additionally, annotation of hydrolytic enzymes was achieved using the CAZy reference sequence database and InterPro protein signature databases. Finally, the fresh supernatants, separated from the cell pellets, were incubated with chitin and then analyzed to identify the size distribution of the chitin degradation products.

**Figure 3:**
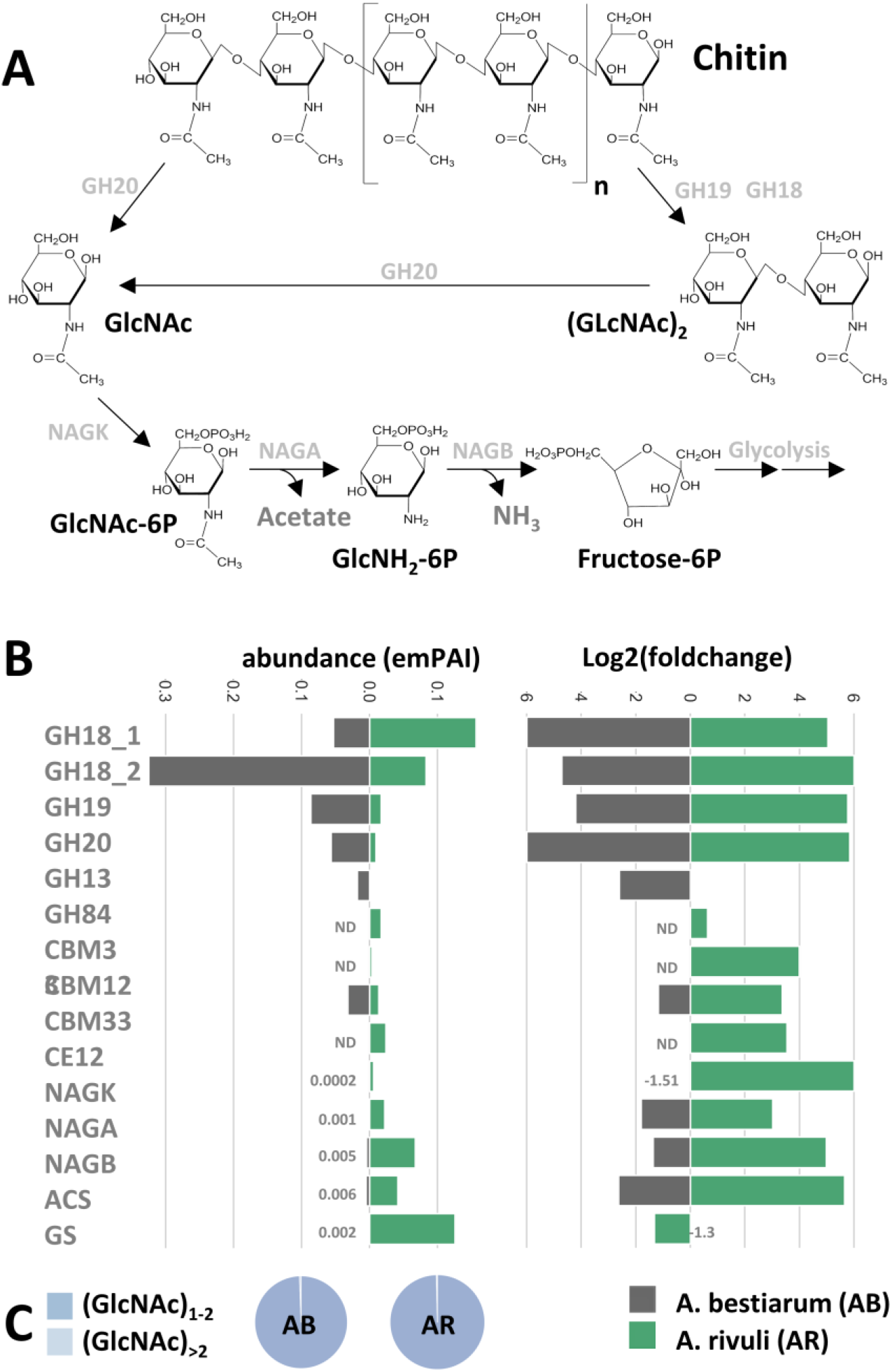
Figure 3: A) The diagram outlines the primary routes of chitin degradation, based on observed enzymes in the secretome and cellular enzymes. First, chitin is cleaved into GlcNAc oligomers by several endochitinases (GH18_1 GH18_2 and GH19), followed by terminal cleavage through exo-chitinase (GH20). After uptake, GlcNAc oligomers are further broken down into monomers and GlcNAc is converted to GlcNAc-6P by NAGK within the cell, and then to GlcNH2-6P by NAGA, before finally transforming into Fructose-6-phosphate (F6P) by NAGB. F6P can then enter glycolysis. The released acetate can also be used to produce acetyl-CoA and ammonia can be incorporated into glutamine. The different potential uptake and transport routes for both *Aeromonas* species are illustrated in Figure 4. B) The bar graphs represent the abundance (emPAI index, when grown on chitin) and log2(foldchange) for the identified glycoside hydrolases and catabolic enzymes in both *Aeromonas* strains when grown on chitin compared to glucose. C) Chitin degradation assays using the secreted enzymes from both cell cultures revealed that primarily GlcNAc monomers and dimers are produced for both strains.

**Figure 4:**
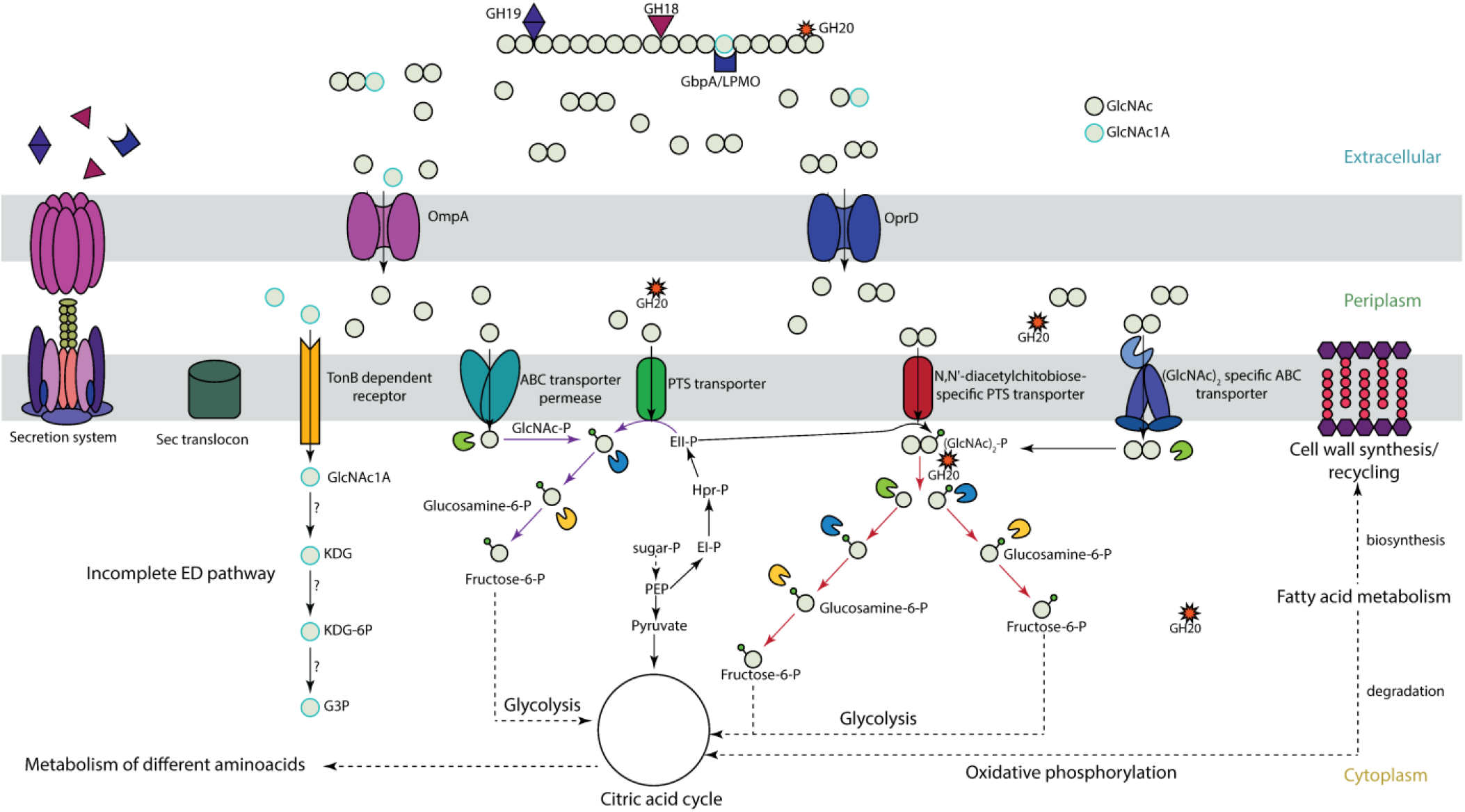
Putative chitin degradation and uptake routes present in *A. bestiarum* and *A. rivuli* based on identified hydrolytic enzymes, transporters, and metabolic enzymes. Chitin is degraded outside the cell by endo- and exo-chitinases (GH18 and GH19) and beta-hexosaminidases (GH20) (see also Figure 3). The hydrolysis products, GlcNAc mono- and oligomers are then transported into the periplasm through dedicated outer membrane porins such OprD, which was highly upregulated when grown on chitin. The chitin degradation assays furthermore demonstrated that mainly GlcNAc mono- and dimers are produced (SI Figure 5). In the periplasm, GlcNAc dimers are further cleaved by N-acetyl hexosaminidases (GH20). Hydrolysis products may then be further transported into the cytoplasm through a N-acetylglucosamine-specific phospho-transferase system (PTS), bind to substrate-binding proteins related to different ABC transporters, or are transported through a dedicated (GlcNAc)_2_ ABC-type transporter. Interestingly, *A. rivuli* possesses an additional (GlcNAc)_2_ specific PTS transporter. Finally, oxidative degradation products (GlcNAc1A) produced through LPMOs are transported inside the cell where it can be directly deacetylated and deaminated to produce KDG without activation by phosphorylation. However, for *Aeromonas* this route could not be confirmed in the present study. Finally, GlcNAc dimers get cleaved by dedicated GH20s and GlcNAc gets further phosphorylated, where GlcNAc-6P is converted into GlcNH_2_-6P and finally into fructose-6-phosphate (F6P), by glucosamine-6-phosphate deaminase, allowing it to enter glycolysis.

### Cellular proteome and secretome response when switching to growth on chitin

When comparing the proteomes of *Aeromonas* grown on glucose with those grown on chitin we observed a significant reorganization of the cellular proteome (Figure 2). For example, in *A. bestiarum*, from 1556 quantified protein groups, 1035 (>65%) showed a fold change of greater than 1.5. In *A. rivuli*, several central metabolic pathways were significantly more abundant when grown on chitin compared to growth on glucose, including the glyoxylate cycle, pentose phosphate pathway, beta oxidation, and degradation of lysine, leucine, histidine, and parts of the TCA cycle. However, glycolytic enzymes (Embden-Meyerhof pathway) and most other biosynthesis pathways were less abundant (SI EXCEL DOC 1 and SI Figure 6). The same was observed for *A. bestiarum*, where the glyoxylate cycle, beta-oxidation, degradation of amino acids, and parts of the TCA cycle were upregulated, but glycolytic enzymes and gluconeogenesis were downregulated (SI EXCEL DOC 1 and SI Figure 6). In addition, several other proteins in the cell biomass were significantly more abundant during growth on chitin, hinting towards their involvement in the degradation, uptake or catabolization of this biopolymer. For *A. rivuli*, 3 different glycoside hydrolases (2 x GH20 and a GH13), one carbohydrate binding protein and 4 different outer membrane porins were strongly upregulated. For example, the porin OprD increased 40 times in abundance in cells grown on chitin. Additionally, 3 PTS transporters (including one N-acetylglucosamine-specific transporter) were strongly upregulated, and a range of different substrate-binding proteins related to different ABC transporters increased in abundance (SI EXCEL DOC 1, Figure 1). The same was observed for *A. bestiarum* where 4 different glycoside hydrolases (2 x GH18 and 2 x GH20), 2 carbohydrate binding proteins and an outer membrane porin (OprD) strongly increased in abundance. Like in *A. rivuli*, OprD was one of the most abundant proteins and it increased over 60 times in abundance in response to chitin. Furthermore, we detected one up-regulated N-acetylglucosamine PTS system and several substrate-binding proteins related to different ABC transporters (SI EXCEL DOC 1, Figure 1, SI Figure 7 and 8). Quantitative comparison of the secretomes from cultures grown on glucose and chitin showed a range of upregulated extracellular hydrolytic enzymes and chitin binding proteins. For example, in *A. rivuli*, 5 chitin-specialized glycoside hydrolases (2 x GH18, GH19, GH20 and GH84), a putative carbohydrate esterase-deacetylase (CE12) and 2 sugar binding proteins (one GlcNAc binding protein, GbpA), a range of outer membrane porins and several ABC transporters were also abundantly present in the secretome of cells which were grown on chitin.

Similarly, the supernatant from *A. bestiarum* revealed 5 different glycoside hydrolases (2 x GH18, 2 x GH20 and one GH13) a GlcNAc binding protein (GbpA), one outer membrane porin, several other proteins related to PTS systems and different solute binding proteins that significantly increased abundance (SI EXCEL DOC 1, Figure 1, SI Figure 7 and 8).

### Chitin degradation and utilization routes in *A. rivuli* and *A. bestiarum*

The secretome and cellular proteome compared between growth on glucose and chitin as well as the oligo- and monomer ratios obtained from the chitin degradation assays allowed to elaborate on the putative chitin degradation routes in both *Aeromonas* strains. Analysis of chitin degradation products revealed domination of GlcNAc dimer and monomers for both strains (SI Figure 5) with only trace amounts of oxidation products (i.e. GlcNAc1A) and deacetylated forms. Consequently, the hydrolysis product profile already indicated that both *Aeromonas* strains predominantly utilize GlcNAc dimer- and monomers (Figure 3c). Furthermore, we identified a similar set of 2 x GH18s (WP_042040344.1, WP_224432263.1, and A0A291U6Z3, A0A291TVA9), one GH19 (WP_084218236.1 and A0A291U6T4) and one GH20 (WP_224432421.1 and A0A291U507) in both *Aeromonas* strains (SI EXCEL DOC 1 and Figure 3b). However, while both GH18s from *A. rivuli* and *A. bestiarum* showed a very high sequence identity (>80%), the GH19s had only a sequence identity of less than 50%, and the GH20s less than 25% (SI EXCEL DOC 2). Family 18 and 19 glycoside hydrolases are endo-acting enzymes that break down chitin at internal sites, forming chitobiose, chitotriose, and chitotetraoses. Family GH20 includes N-acetyl-glucosaminidases which act on non-reducing ends to either release dimers (chitobiose) or to further break down multimer products into GlcNAc (25). The GH18 and GH19 families do not share sequence similarity. GH18 chitinases cleave the chitin into β-anomer products, whereas GH19 hydrolyze chitin to α-anomer by using the inverting mechanism (26, 27). Interestingly, both strains express 2 different GH18 chitinases, one with an approx. MW of 90 kDa and a second with a MW of approx. 105 kDa. Analysis of the amino acid sequence against the InterPro databases showed that the smaller GH18 chitinase (*A. rivuli*, WP_224432263.1, 90 kDa) includes next to the glycoside hydrolases family 18 domain (PF00704) also a Chitinase A N-terminal domain (PF08329), a PKD/REJ-like domain (PF02010) and two carbohydrate-binding modules family 5/12 (IPR003610, also annotated with PF02839). The latter (PF02839) is known to specifically bind to insoluble chitin (28). The amino acid sequence of the larger GH18 (A. rivuli WP_042040344, 105 kDa) contained next to the glycoside hydrolases family 18 domain (PF00704) a Chitinase C domain (PF06483) and a bacterial Ig domain (PF17957), and two carbohydrate-binding family 5/12 modules (IPR003610). Nevertheless earlier studies on the different chitinases (ChiA, ChiB, ChiC, ChiD, ChiE, ChiF, ChiG, and ChiH) demonstrated that these show different hydrolytic activities against the different forms of chitin (29, 30). Intriguingly, in *A. rivuli* the GH18 Chitinase C was higher expressed, while in *A. bestiarum* the GH18 Chitinase A was more abundant (Figure 3B, GH18_1 Chitinase A domain; GH18_2 Chitinase C domain). Additionally, two others, but lower-abundance glycoside hydrolases (GH13 and GH84) were observed in either *A. rivuli*, or *A*. bestiarum growing on chitin. While GH13 family enzymes are specialized on alpha-glucans, the detected GH84 (WP_224432541) contains a beta-N-acetylglucosaminidase catalytic domain. Yet, the presence of GH18, 19 and 20 hydrolases seems to be sufficient to explain effective growth of the studied species on insoluble chitin. Furthermore, their abundance differences between both strains explains the observed ratio differences between the GlcNAc mono- and dimers in the chitin degradation experiment. Also, one carbohydrate binding protein that was detected in the supernatant of both strains was classified as CAZy family AA10 (formerly CBM33), which is a copper-dependent lytic polysaccharide monooxygenase (LPMO) (SI EXCEL DOC 2 and 3). These enzymes catalyze the cleavage of 1,4-glycosidic bonds found in different types of plant cell wall polysaccharides and chitin. LPMOs function on the crystalline regions of polysaccharides, thereby making crystalline chitin better available to other hydrolytic enzymes. The LMPOs found in *A. rivuli* and *A. bestiarum* showed a sequence identity of >70% and significantly increased in abundance during growth on chitin (SI EXCEL DOC 2). However, the changes in protein level were much less pronounced compared to the changes observed for the other chitinases (SI EXCEL DOC1). One possible explanation for this is that the growth experiments utilized amorphous chitin, eliminating the need for accelerating degradation of crystalline chitin. In agreement with this, only trace quantities of GlcNAc1A were detected in the chitin hydrolysis assays (SI Figure 5). Interestingly, N-acetyl-hexosaminidases (GH20) have been reported to be located mainly in the periplasm. However, in particular in the secretome of *A. bestiarum* we detected larger quantities of a GH20 (A0A291U507, SI EXCEL DOC 1), which according to SignalP prediction (SignalP-6.0) however contains a Sec/SPI signal, which could allow to cross the outer membrane in Gram-negative bacteria. Previous studies have shown that chitin monomers and dimers uptake can be enhanced through outer membrane porins (Figure 4). Specifically, Kitaoku *et. al*., reported on a chitoporin specific to chitin found in *Vibrio spp* (31). Homologues can also be found in other proteomes, such as *E. coli* and *A. veronii*. To further investigate this, we performed BLAST search on the genomes of *A. bestiarum* and *A. rivuli* using the reported chitoporins from *A. veronii* and *E. coli* (SI EXCEL DOC 2). Thereby, we found a high sequence identity with the OprD family outer membrane porins in both *Aeromonas* strains (*A. rivuli* WP_042040660 and *A. bestiarum* A0A291U719), which were also highly upregulated during growth on chitin. Therefore, it can be hypothesized that the OprDs identified in both strains are chitoporins which further support the uptake of GlcNAc monomers and dimers into the periplasm. The GlcNAc dimers can then be further cleaved by periplasmatic GH20s into monomers. In fact, we found abundant GH20s in the cellular proteomics experiments of both strains (*A. rivuli* WP_042039723 and *A. bestiarum* A0A291U507, SI EXCEL DOC 1). Different mechanism have been reported which transport monomers and dimers through the inner cell membrane (Figure 4) (9, 14, 31, 32).

**Figure 5:**
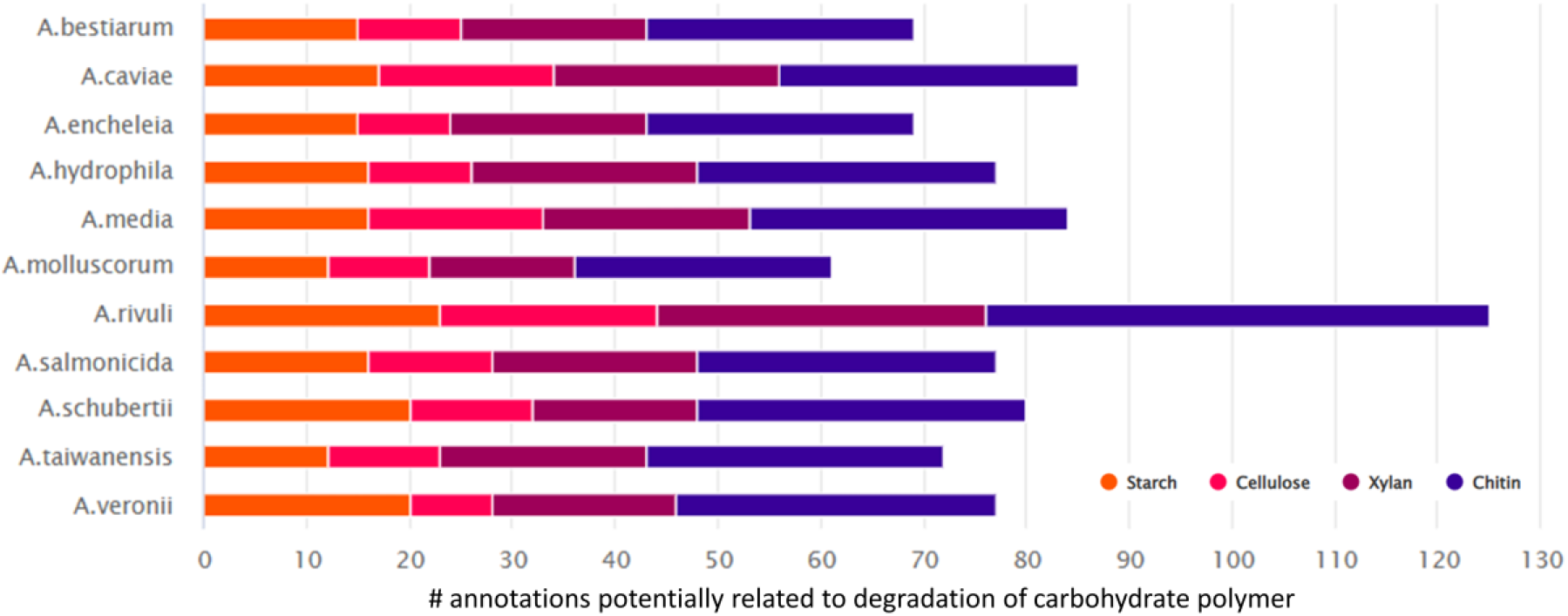
The bar graph displays the frequency of genes associated with CAZy families involved in hydrolyzing starch, cellulose, xylan, and chitin. The analysis concentrated on *Aeromonas* strains, which are commonly found in drinking water distribution systems. A large spectrum of glycoside hydrolases (GHses) related to the degradation of starch, cellulose, xylan and chitin were identified in the genomes of the selected *Aeromnas* strains (SI DOC TABLE 1).

In fact, several substrate-binding domain-containing proteins related to different ABC transporters (which potentially transport the GlcNAc monomers and dimers) were found to be upregulated when grown on chitin (SI EXCEL DOC 1). GlcNAc monomers are then directly phosphorylated, while the dimers are cleaved and phosphorylated by a GlcNAc specific kinase (*A. rivuli* WP_042041492, *A. bestiarum* A0A291U705), which then releases GlcNAc and GlcNAcP (Figure 4). Moreover, we also identified N-acetylglucosamine-specific phosphotransferase system (PTS) which was upregulated during growth on chitin (*A. rivuli* WP_042039724 and *A. bestiarum* A0A291U5N4, which was annotated as ‘glucose specific’ albeit having 96% sequence identify with WP_042039724), this transporter simultaneously phosphorylates and transports GlcNAc monomers into the cytoplasm. Li *et al*., (2007) reported on a putative (GlcNAc)2 catabolic operon in *V. cholerae* (11). This organism is from the same Gammaproteobacteria class as the here studied *Aeromonas* strains. This operon includes next to the chitin sensor *chiS* also the chitin binding protein ‘(GlcNAc)2 periplasmic binding protein’ and a related ABC-type (GlcNAc)2 transporter (33). The periplasmic binding protein binds chitin oligomers (e.g. GlcNAc dimers) upon which it dissociates from *chiS*. This triggers expression of the chitinolytic genes. BLAST search confirmed that homologues genes (with high sequence similarity) are also present in both investigated *Aeromonas* strains (SI EXCEL DOC 2). Furthermore, the putative (GlcNAc)2 periplasmic binding protein (*A. rivuli* WP_042041499.1 and *A. bestiarum* A0A291U0Y6) of the ABC-type (GlcNAc)2 transporter was strongly upregulated and abundant during growth on chitin (SI EXCEL DOC 1).

Finally, a putative diacetyl-chitobiose specific PTS was detected in the genome of *A. rivuli*. This system facilitates the phosphorylation of GlcNAc dimers alongside their translocation across the inner membrane. However, a homologous gene could not be identified in the genome of *A. bestiarum*. Such PTS transporters have previously been described for other Gammaproteobacteria such as *S. marcescens* or *E. coli* (34, 35). However, the known subunits (EIIA WP_224432960.1, EIIB WP_224432412.1 and EIIC WP_042041548.1) were barely detected and upregulation in response to chitin could therefore not be confirmed. In the cytoplasm, GlcNAc dimers are then supposedly cleaved by another GH20. Indeed, the cellular proteomics experiments revealed a strongly upregulated GH20 (*A. rivuli* WP_224432421.1, *A. bestiarum* A0A291U0V8) which does not contain a signal peptide (and which therefore likely resides in the cytoplasm, SI EXCEL DOC 1). The GlcNAc monomers are then further converted into glucosamine-6-phosphate by N-acetylglucosamine kinase (NAGK, for *A. rivuli* either WP_224430983.1 or WP_042040690.1, *A. bestiarum* A0A291TVT4), then into glucosamine-6-phosphate by N-acetylglucosamine-6-phosphate deacetylase (NAGA, *A. rivuli* WP_042039726.1, *A. bestiarum* A0A291U5J9) and finally into fructose-6-phospate (F6P) by Glucosamine-6-phosphate deaminase (NAGB, *A. rivuli* WP_042039725.1, *A. bestiarum* A0A291U4P9) which can enter glycolysis, and the resulting pyruvate (in form of acetyl-CoA) the TCA cycle (Figure 4). Except for NagK in *A. bestiarum*, the enzymes from this pathway were upregulated during growth on chitin.

Noteworthy, during the intracellular conversation of GlcNAc to glucosamine, acetate is also released which can be further converted to acetyl-CoA by the acetyl-CoA synthetase. In fact, the acetyl-CoA synthetase/acetate-CoA ligase was overexpressed in *A. bestiarum* and *A. rivuli* when grown on chitin (*A. rivuli* WP_224432291.1, *A. bestiarum* A0A291U0K7). Another pathway that can utilize acetate is the glyoxylate cycle, which was found to be significantly overexpressed in both *Aeromonas* strains. Chitin furthermore can be an important nitrogen source in environments where nitrate and ammonia concentrations are low, such as in drinking water systems.

Glucosamine-6-phosphate deaminase (NAGB) releases ammonia which can be assimilated through incorporation into glutamine, a proteogenic amino acid and an important nitrogen carrier (36). This reaction is facilitated by glutamine synthase (GS), which however was only overexpressed in *A. bestiarum* (*A. rivuli* WP_042042627.1, *A. bestiarum* A0A291TY36). Finally, we wanted to explore whether both *Aeromonas* strains have the potential to utilize oxidative degradation products (GlcNAc1A) as recently proposed by Jiang *et al*. (2022)(9). The oxidative degradation of chitin through LPMOs accelerates the bioconversion of crystalline chitin. Jiang *et al*., demonstrated that GlcNAc1A can be directly deacetylated and deaminated to produce KDG without activation by phosphorylation. KDG is also the precursor of a key intermediate of the Entner-Doudoroff pathway (KDG-6P), which pathway however was found to be incomplete in the genome of both *Aeromonas* strains (SI EXCEL DOC 4). Nevertheless, the utilization of oxidation products through this route is believed to be common in marine Gammaproteobacteria. Albeit *A. bestiarum* and *A. rivuli* have been isolated from fresh waters, other members of the genus *Aeromonas* are widely distributed in estuarine, and marine environments. Therefore, we searched for genes involved in degradation and catabolism of GlcNAc1A (LPMOs, OngA/B/C, KdgK and KdgA) as described by Sheng *et al*., (9) in the genomes of *A. bestiarum* and *A. rivuli*. Interestingly, while homologues for LPMOs (*A. rivuli* WP_224431348.1, *A. bestiarum* A0A291TWD8), and KdgK (*A. rivuli* WP_042040847.1, *A. bestiarum* A0A291U805) and KdgA (*A. rivuli* A. WP_224431468.1, *A. bestiarum* A0A291TWZ9) were found, only more distant or no significant alignments were obtained for the key genes OngC or OngB (SI EXCEL DOC 2). While both putative LPMOs were found to be expressed and slightly upregulated when grown on (amorphous) chitin, the putative KdgK and KdgA homologues were not detected in the proteomics experiments. Nevertheless, at present, it cannot be excluded that other deacetylases and deaminases expressed by *Aeromonas* facilitate the conversion of GlcNAc1A.

### The potential to degrade a broader spectrum of carbohydrate biopolymers

The conducted growth and proteomics experiments clearly demonstrate that both *Aeromonas* strains have the ability to efficiently degrade and grow on chitin as the sole carbon and nitrogen source. However, in highly oligotrophic environments such as drinking water distribution systems, the survival of *Aeromonas* may be further enhanced by the ability to degrade, uptake, and utilize of a wider range of different biopolymers.

In order to investigate this, we analyzed the genomes of several additional *Aeromonas* species commonly found in drinking water distribution systems, for dedicated CAZy genes. This revealed a range of genes which are potentially involved in the degradation of different carbohydrate polymers such as starch, cellulose or xylan (Figure 5, Supplementary Information Figure 9). However, it is important to note that the in silico identification of glycoside hydrolases (and related binding proteins) is only a potential, and their actual ability to hydrolyze and utilize specific polysaccharides requires further cultivation experiments.

## Discussions

*Aeromonads* are frequently found in the sediments and loose deposits of drinking water distribution systems (1, 4, 5, 37-40). These deposits contain a variety of organic and inorganic suspended solids, and harbor microscopic fungi, as well as small and larger invertebrates, such as *A. aquaticus* (41-44). These organisms are known to produce biopolymers, including chitin, which can serve as a source of carbon and nitrogen for microbes. The ability to use chitin as a nutrient source is commonly found among bacteria (7-9). However, its specific role within environments such as drinking water distribution systems is not yet fully understood.

The growth experiments conducted here demonstrate that two frequently found *Aeromonas* strains in drinking water distribution systems, *A. bestiarum* and *A. rivuli*, efficiently degrade and utilize chitin as their sole carbon and nitrogen source. This finding demonstrates that *Aeromonas* is not only chitinolytic, but also chitinotrophic. Previous studies have not shown significant growth for *A. rivuli* (and several other strains) in sterilized drinking water and low concentrations of chitin (1). In our study, *A. rivuli* also showed a slower growth compared to *A. bestiarum*, despite the use of a more soluble amorphous chitin and potentially higher substrate concentrations. Our quantitative proteomics analysis of the secretome and cell biomass further revealed that chitin oligomers not only induced the expression of dedicated glycoside hydrolases, sugar binding proteins, and transporters, but also showed strong differences in the global cellular proteome (see Figure 1). This is likely because GlcNAc has to be converted into fructose 1,6-bisphosphate before it can enter glycolysis, and additional acetate and ammonia are released which also provides energy, organic carbon and a nitrogen source (25). Consequently, we identified a dedicated network of chitinases (2x GH18s and one GH19), β-N-acetyl hexosaminidases (GH20) and binding proteins. One GH18 contains a Chitinase A and the second a Chitinase C domain and a carbohydrate-binding module (PF02839) which can bind to insoluble chitin, which was particularly abundant in *A. bestiarum. Aeromonas* species also expressed a GH19, which are commonly only found in plants and some actinobacteria. The few known bacterial GH19s are supposedly acquired from plants through horizontal gene transfer (45). The GH18s in *A. rivuli* and *A. bestiarum* show a high sequence identify, but the GH19 and GH20s appeared to be only distantly related. Nevertheless, the supernatants from both strains efficiently degraded chitin into mono and dimers. GlcNAc dimers and oligomers may provide a competitive advantage since the GlcNAc monomers can be easily utilized by other microorganisms present in the community. Nevertheless, there are also some microbes such as *E*.*coli* which – albeit not being able to hydrolyze chitin – possess transporters and enzymes dedicated to utilization of GlcNAc dimers (chitobiose) (46).

Furthermore, both strains strongly overexpressed a putative chitin specific outer membrane porin (OprD) in response to chitin, which apparently supports the transport of GlcNAc oligomers and monomers through the outer membrane. Membrane transporters are commonly challenging to study because these proteins often show a very hydrophobic nature, are difficult to solubilize, digest and consequently detect. Nevertheless, our study detected several substrate-binding proteins related to different ABC and PTS transporters. Based on these transporters, the uptake mechanisms are diverse and supposedly slightly differ between both strains. For example, *A. rivuli* possesses an additional chitobiose specific PTS sugar transporter, which is not present in the genome of *A. bestiarum*. The recently reported oxidative pathway opens another route to also utilize the oxidation products (GlcNAc1A) generated by LPMOs. However, albeit homologues to LPMOs and other key metabolic enzymes of this pathway are present in both strains, the amorphous chitin used in this study apparently did not lead to relevant amounts of GlcNAc1A. Nevertheless, initial growth experiments on shrimp chitin flakes, which show a high degree of crystallization, did not result in an immediate growth of *A. bestiarum* either. Therefore, the activity of the identified putative LPMOs – which support the degradation of crystalline chitin in other marine microbes (9) – could not be confirmed in this study. Furthermore, both strains secreted also several lipases, peptidases and proteases, which commonly help to make nutrient sources accessible (47, 48). This is also in agreement with the previous finding that feeding Gram-negative bacteria complex sugars can increases the production of lipases (49), which may aid in biofilm formation and play a role in virulence (50, 51). A high growth affinity of *Aeromonas* for long-chain fatty acids in drinking water has been also demonstrated earlier (2).

Nevertheless, albeit *Aeromonas* can efficiently degrade and grow on chitin, likely also several other microbes contribute to the degradation of chitin and other biopolymers in such environments (52, 53). For example, earlier studies also showed that *Aeromonas* is only a minor part of the microbial community present in the drinking water distribution system (4, 54), and other microbes could make crystalline chitin more accessible within the food web. The metabolic end products may furthermore support other heterotrophic bacteria in the drinking water environment. Such interactions have been reported for other niches, such as the soil environments, recently (55-57).

In summary, this study demonstrates how *Aeromonas* can grow on the carbohydrate polymer chitin available in the biomass of invertebrates such as *A. aquaticus*, often found in the loose deposits in drinking water distribution system. Additionally, the quantitative proteomics data reveals a dedicated chitin degradation and uptake network, providing a valuable resource for further investigation of the identified hydrolytic enzymes, transporters, and catabolic enzymes. A deeper understanding of the metabolic routes in these microbes supports the development of better water sanitation strategies.

## Supporting information

SI DOC

SI EXCEL DOC 1

SI EXCEL DOC 2

SI EXCEL DOC 3

SI EXCEL DOC 4

## Acknowledgements

The authors would like to thank Dita Heikens for her support in the proteomics laboratory, and all colleagues in the Department of Biotechnology, as well as Professor Bert van der Wal, for valuable discussions. They would also like to acknowledge Evides (The water company N.V.) and the NWO Spinoza prize awarded to Mark van Loosdrecht for funding and support.

