## Supplementary material for "Exploring the metabolic potential of *Aeromonas* to utilise the carbohydrate polymer chitin": SI DOC

**TABLE OF CONTENTS** **PAGE**

SI Figure 1: OD660 of *A. bestiarum* and *A. rivuli* cultures grown on glucose and chitin. 2

SI Figure 2: Microscopy images of the *A. bestiarum* and *A. rivuli* grown on glucose and chitin. 3

Si Figure 3: PCA analysis of replicate proteome profiles for *A. bestiarum* and *A. rivuli*. 4

SI Figure 4: Hierarchical clustering of replicate proteome profiles for *A. bestiarum* and *A. rivuli*. 5

SI Figure 5: Chitin degradation assay with A. bestiarum and A. rivuli cell culture supernatants. 6

SI Figure 6: Changes in CCM in A. bestiarum and A. rivuli between growth on glucose and chitin. 7

SI Figure 7: Changes in CAZy expression profiles in A. bestiarum between growth on glucose and chitin. 8

SI Figure 8: Changes in CAZy expression profiles in A. rivuli between growth on glucose and chitin. 9

SI Figure 9: CAZy and binding-proteins involved in degradation of chitin for different Aeromonas strains. 10

SI Table1: CAZy families potentially involved in the degradation of different biopolymers. 11

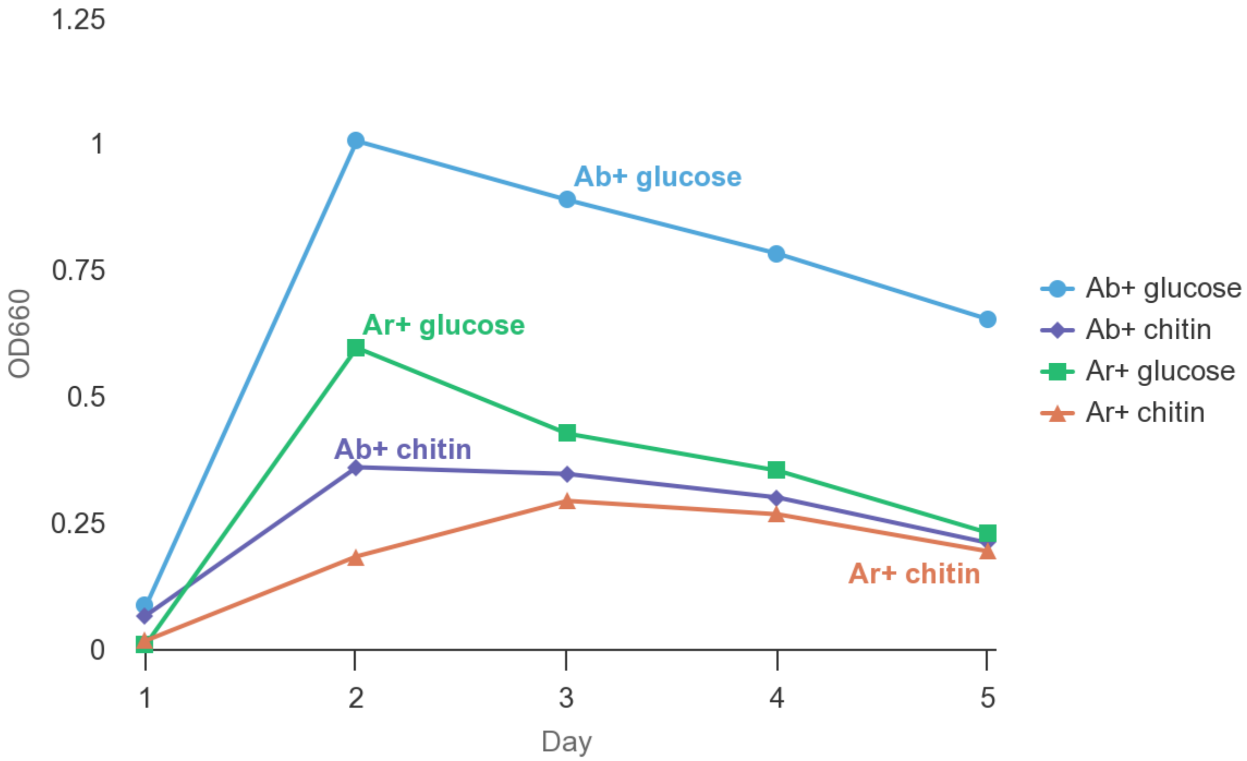


**SI Figure 1.** OD660 measurements for *A. bestiarum* (Ab) and *A. rivuli* (Ar) cultures grown on glucose or chitin over 5 days.

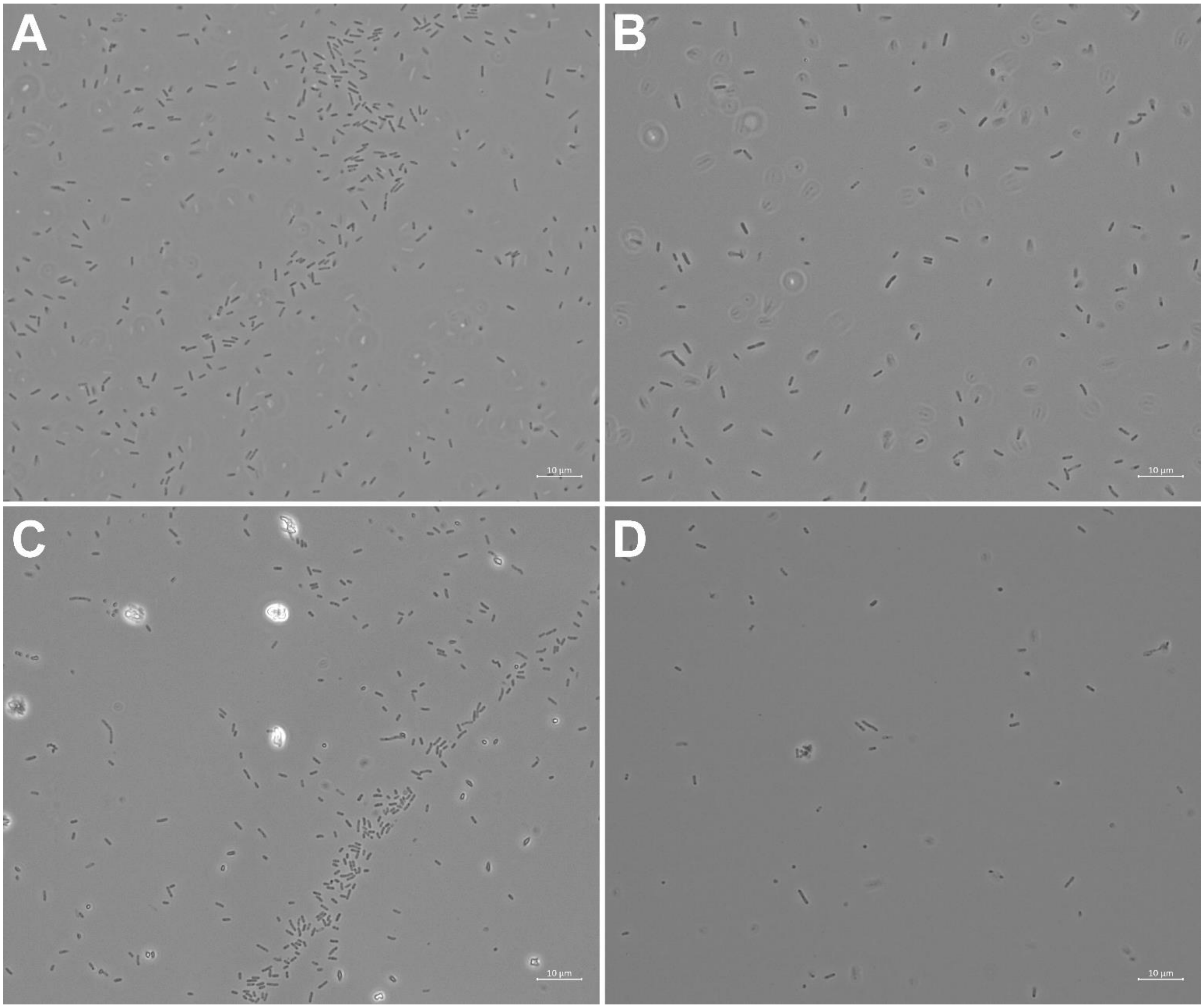


**SI Figure 2.** Light microscopy images (at 100x magnification) of cultures from (A) *A. bestiarum* grown on chitin, (B) *A. bestiarum* grown on glucose, (C) *A. rivuli* grown on glucose, and (D) *A. rivuli* grown on chitin.

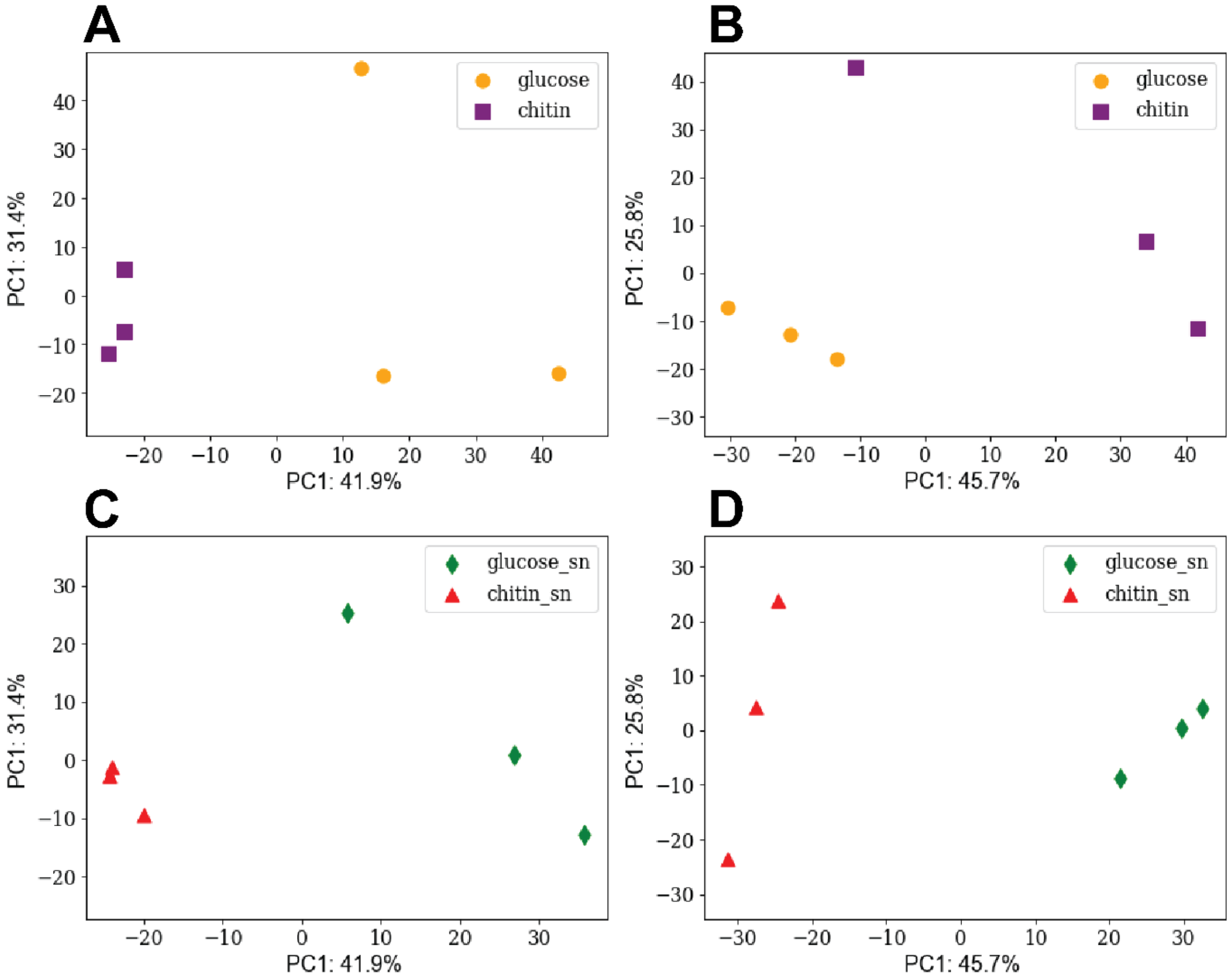


**SI Figure 3**: The graphs display the principal component analysis (PCA) of the profiles acquired from the triplicate growth experiments of (A) *A. bestiarum* biomass, (B) *A. rivuli* biomass, (C) *A. bestiarum* secretome, and (D) *A. rivuli* secretome. Supernatant is abbreviated as "sn".

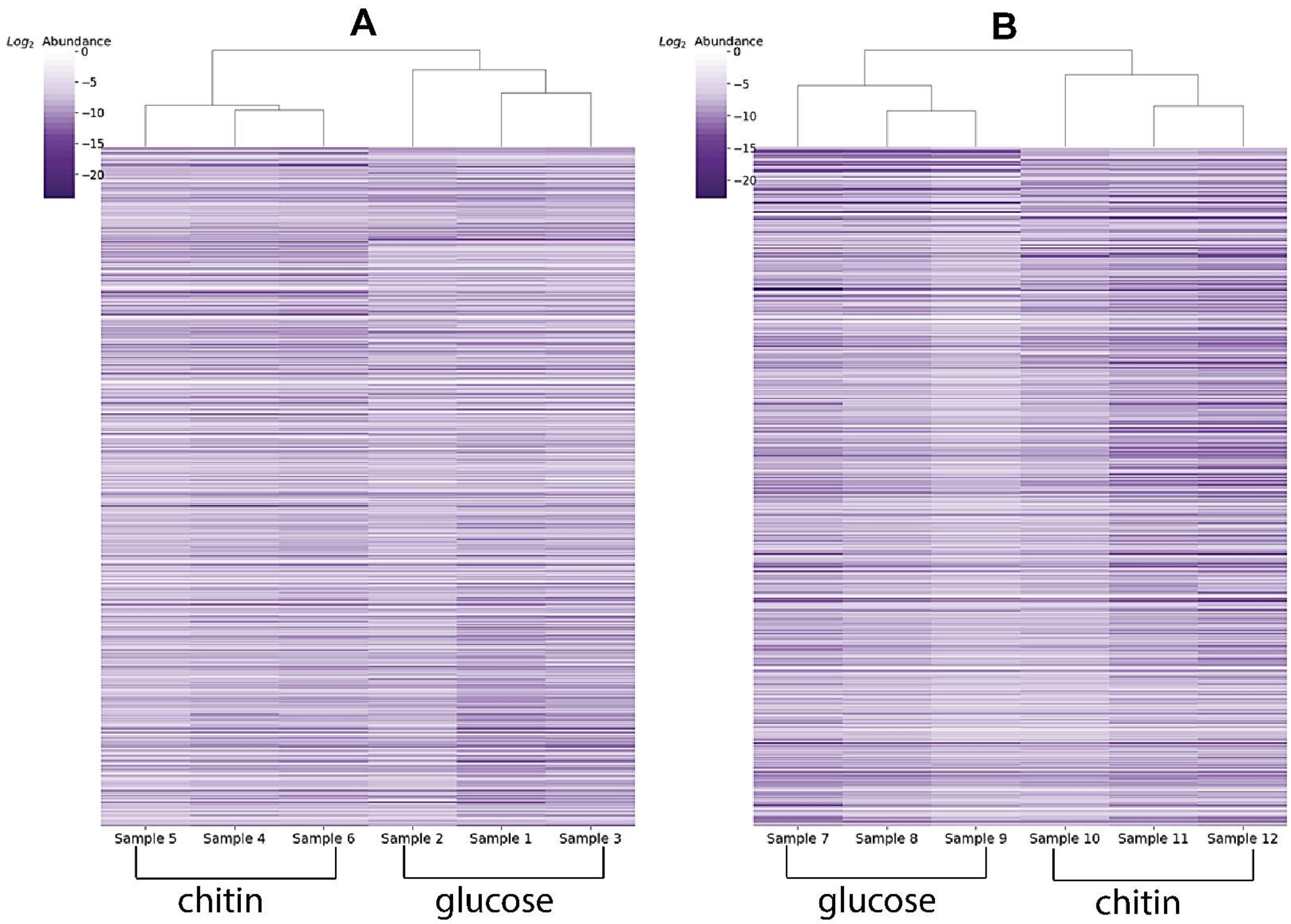


**SI Figure 4:** The heatmaps display the hierarchical clustering of the cellular proteome profiles acquired from the triplicate growth experiments of (A) *A. bestiarum*, and (B) *A. rivuli*.

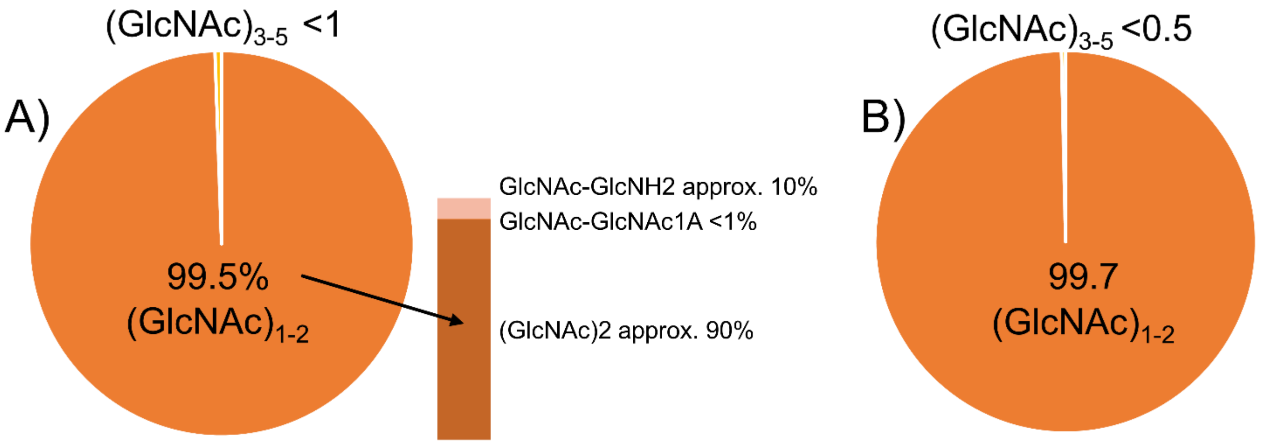


**SI Figure 5:** The pie charts display the distribution between different GlcNAc hydrolysis products (monomers = $(GlcNAc)_1$, dimers = $(GlcNAc)_2$, trimers = $(GlcNAc)_3$, and tetramers = $(GlcNAc)_4$) obtained by exposing chitin to the cell culture supernatants of A) *A. bestiarum* and B) *A. rivuli*. The abundances are based on the different fragment ion intensities of the respective hydrolysis products. Interestingly, the ratios $(GlcNAc)_2$ to $(GlcNAc)_1$ is inverted between both Aeromonas species. For both strains small quantities of oxidized forms could be detected. However, all were <1% in abundance compared to the main hydrolysis product. For *A. bestiarum* partially deacetylated forms could also be detected (approx. 10%). The m/z values for the native hydrolysis products are: GlcNAc = 222.09721, C8H16NO6+; GlcNAc-GlcNAc = C16H29N2O11+, 425.17659; and for the oxidized forms are: GlcNAc1A = 238.09213, C8H16NO7+; GlcNAc-GlcNAc1A = C16H29N2O12+, 441.1715, and for the native deacetylated forms are: GlcNAc-GlcNH2 = 383.16602, C14H27N2O10+.

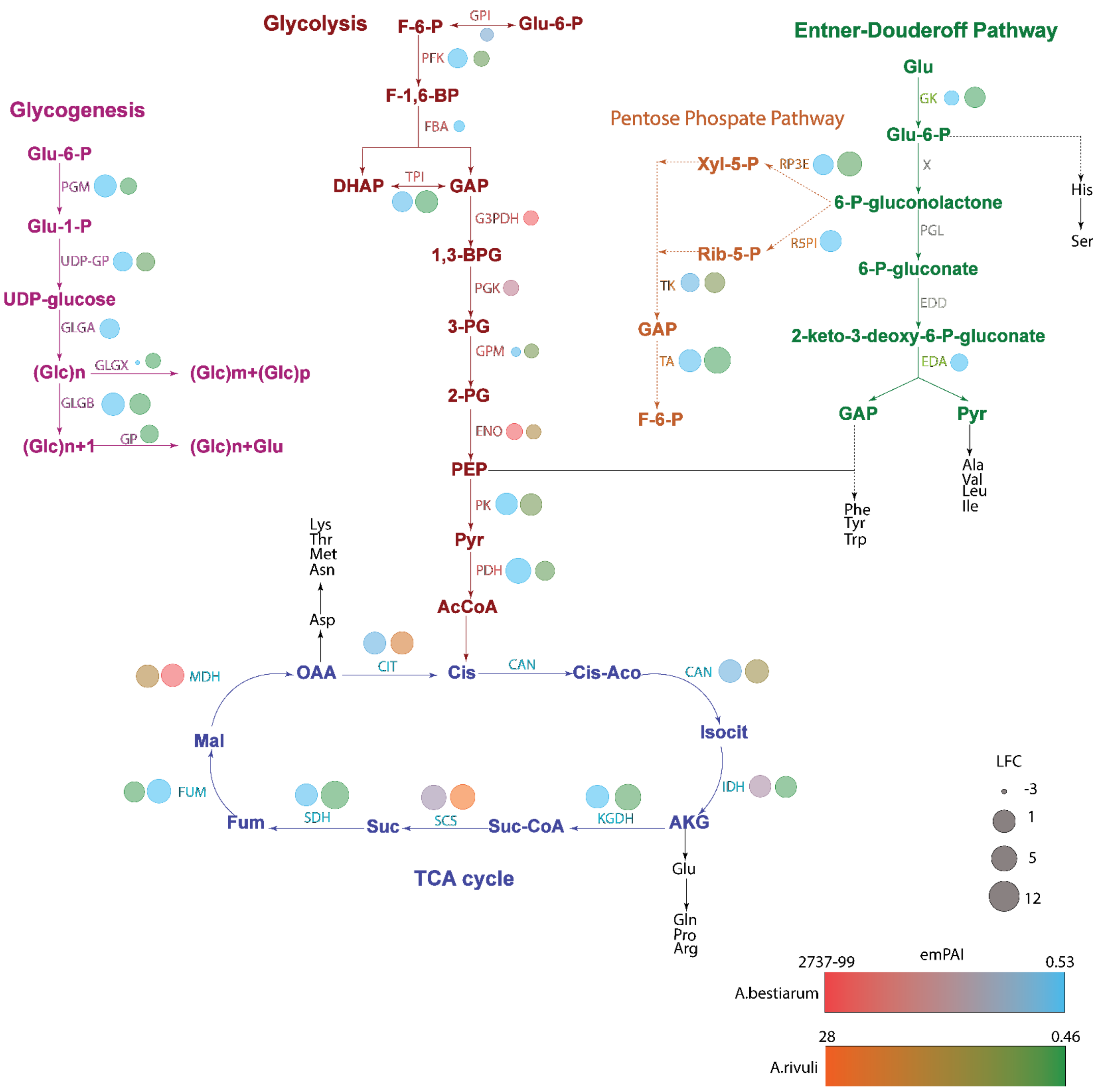


**SI Figure 6:** The graph illustrates abundances of enzymes of the central carbon metabolism and fold changes between growth on glucose and chitin, for *A. bestiarum* and *A. rivuli*. The abundance of the enzymes is represented by the emPAI indices and respective color gradients. The log2(fold change) between growth on glucose and chitin correlates with the circle size. The left circles represent the abundance of enzymes in *A. bestiarum*, and the right circles represent the abundance of enzymes in *A. rivuli.* Enzymes that were not observed in the proteomics experiments are kept in grey (e.g. "PGL" and "EDD"). The genomes of *A. bestiarum* and *A. rivuli* encode only for an incomplete ED pathway (SI EXCEL DOC 4).

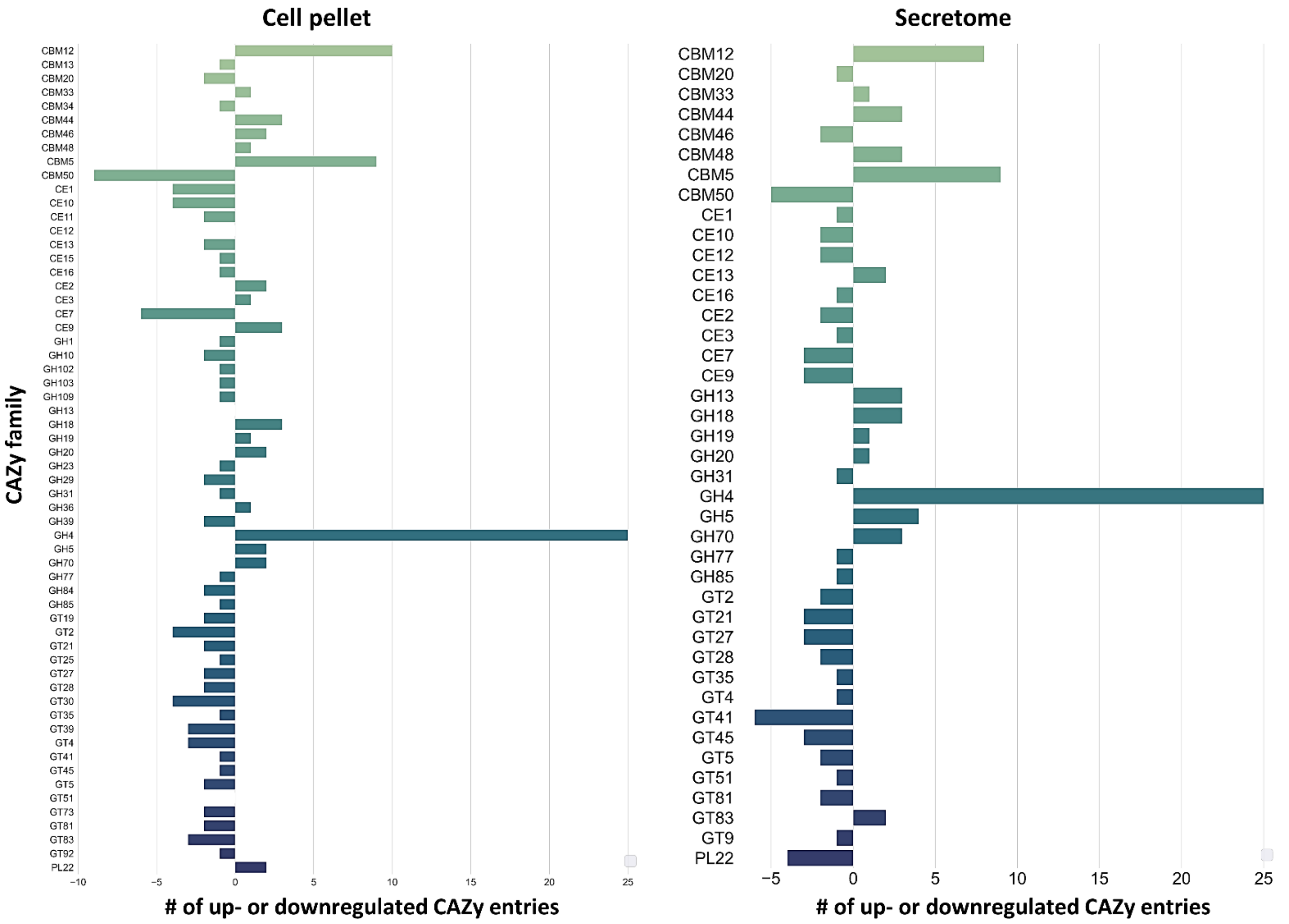


**SI Figure 7**: The bar graph shows the up- or downregulated CAZy families (fold change >1, or <1) for *A. bestiarum*, when switching from glucose to chitin as carbon source. The size of the bars correlates with the number of proteins from this CAZy family. The left graph shows the changes observed in the cellular proteomics experiment, where the right graph shows the changes observed in the secretome analysis.

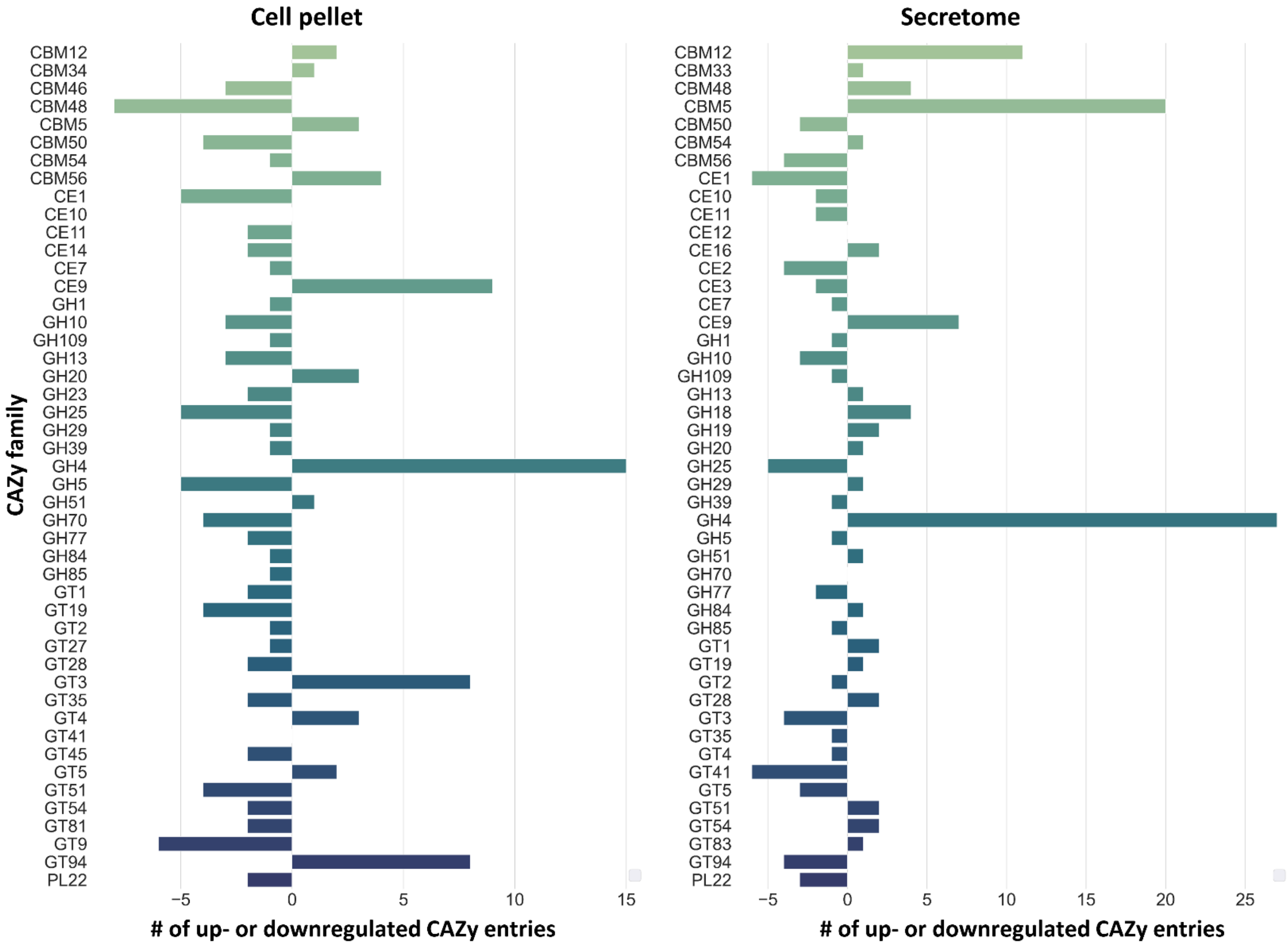


**SI Figure 8:** The bar graph shows the up- or downregulated CAZy families (fold change >1, or <1) for *A. rivuli*, when switching from glucose to chitin as carbon source. The size of the bars correlates with the number of proteins from this CAZy family. The left graph shows the changes observed in the cellular proteomics experiment, where the right graph shows the changes observed in the secretome analysis.

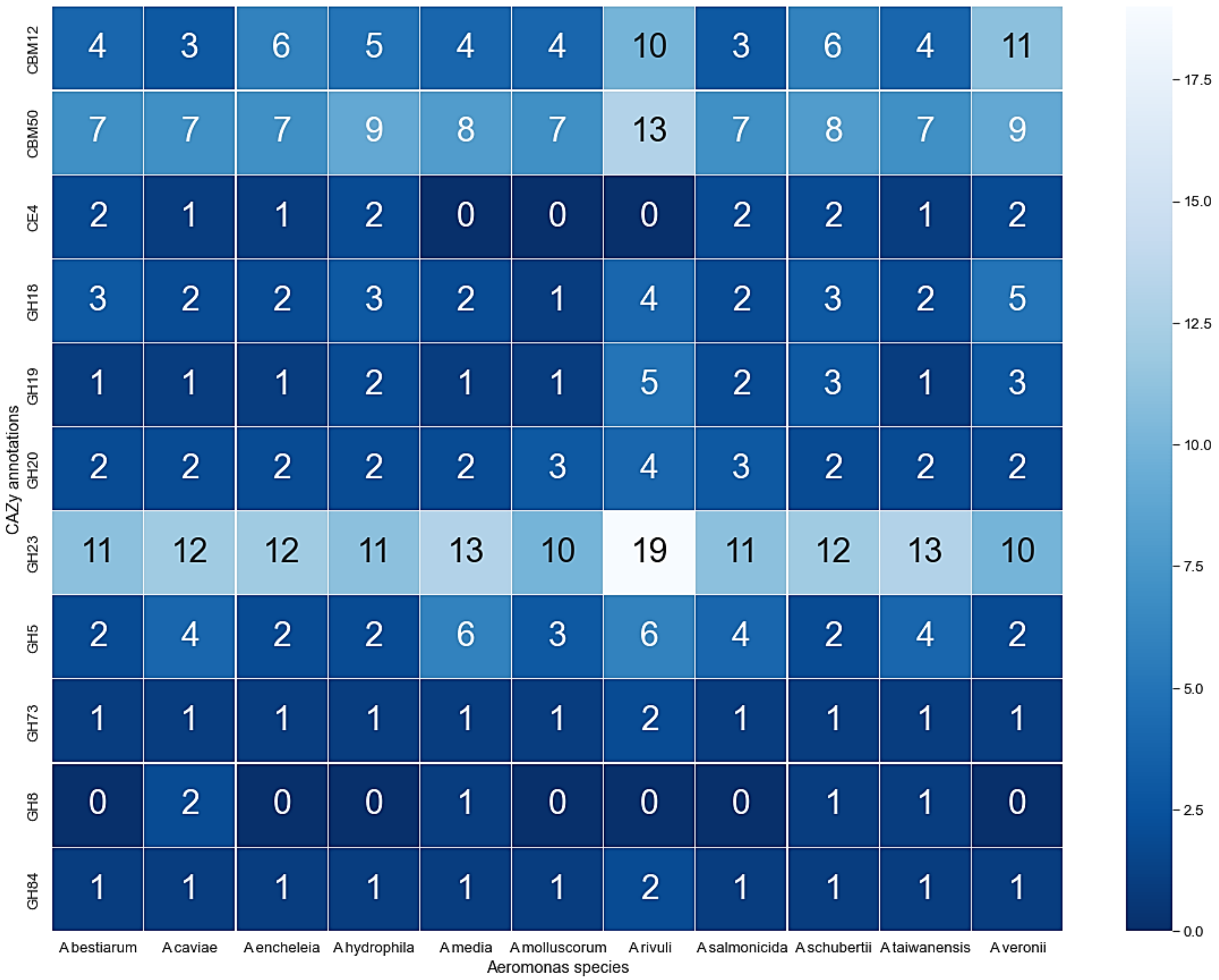


**SI Figure 9:** The heatmap shows the number of identified glycoside hydrolases (GH), related carbohydrate-binding modules (CBM) and chitin esterases (CE) in the genomes of different Aeromonas species. These are potentially involved in the breakdown of chitin and chitosan. The analyzed species are frequently found in drinking water distribution systems.

SI Table 1: The table lists CAZy database families used to search for the potential to degrade various carbohydrate biopolymers in Aeromonas genomes.

| **Biopolymer** | **CAZy targets** |
|---|---|
| Chitin | CBM12, CBM14, CBM18, CBM50, CBM55, GH18, GH19, GH23, GH48, CBM32, GH18, GH20, GH73, GH84, GH85, GH89, GH111, GH116, GH163, GH5, GH7, GH8, GH46, GH75, GH80, CE4 |
| Xylan | AA10, AA14, CBM2, CBM4, CBM6, CBM9, CBM13, CBM15, CBM22, CBM31, CBM35, CBM36, CBM42, CBM54, CBM59, CBM60, CBM72, CBM91, CE1, CE2, CE3, CE4, CE5, CE6, CE7, CE12, GH3, GH5, GH8, GH10, GH11, GH18, GH26, GH30, GH43, GH51, GH67, GH98, GH115, GH141, GT8, GT43, GT47, GT61 |
| Cellulose | AA9, AA10, AA15, AA16, CBM1, CBM2, CBM3, CBM4, CBM6, CBM8, CBM9, CBM10, CBM16, CBM17, CBM28, CBM30, CBM37, CBM44, CBM46, CBM49, CBM59, CBM63, CBM64, CBM72, GH5, GH8, GT2 |
| Starch | AA13, CBM20, CBM21, CBM25, CBM26, CBM34, CBM45, CBM53, CBM69, CBM74, CBM82, CBM83, GT5, GT35, GH13, GH14, GH57, GH126, GH15, GH57, GH97, GH119 |
| Chitosan | GH3, GH5, GH7, GH8, GH18, GH46, GH75, GH80 |
